# Proton exchange in the nitrate vacuolar transporter AtCLCa is required for growth and nitrogen use efficiency

**DOI:** 10.1101/2022.09.21.508937

**Authors:** Julie Hodin, Christof Lind, Anne Marmagne, Christelle Espagne, Michele Wolfe Bianchi, Alexis De Angeli, Fadi Abou-Choucha, Mickaël Bourge, Fabien Chardon, Sebastien Thomine, Sophie Filleur

**Affiliations:** Université Paris-Saclay, CEA, CNRS, Institute for Integrative Biology of the Cell (I2BC), 91198, Gif-sur- Yvette, France; Université Paris Cité, UFR Sciences du Vivant, 35 rue Hélène Brion, F-75205 Paris Cedex 13, France; Université Paris-Saclay, INRAE, AgroParisTech, Institut Jean-Pierre Bourgin (IJPB), 78000, Versailles, France; Université Paris-Est-Créteil-Val-de-Marne, 61 avenue du Général de Gaulle, 94010 Créteil Cedex France

## Abstract

Nitrate is a major nutrient and osmoticum for plants. To deal with its fluctuating availability in soils, plants store it into vacuoles. AtCLCa, a 2NO_3_^-^/1H^+^ exchanger localized on the vacuole ensures this storage process. It belongs to the CLC family that includes exchangers and channels. A mutation in a glutamate residue conserved across CLC exchangers is likely responsible for the conversion of exchangers to channels. Here, we show that a *clca* mutant of this residue, E203, behaves as an anion channel in its native membrane. To investigate its physiological importance, we introduced the *AtCLCa_E203A_*point mutation in a *clca* KO mutant. We first showed that these *AtCLCa_E203A_* mutants display a growth deficit linked to water homeostasis disruption. Additionally, *AtCLCa_E203A_*expression is not able to complement the *clca* defect in nitrate accumulation and favors higher N-assimilation at the vegetative stage. Further analyses at post-flowering stages indicated that AtCLCa_E203A_ results in an increase of N uptake allocation to seeds, leading to a higher nitrogen use efficiency compared to wild-type. Altogether, these results point out the critical function of the AtCLCa exchanger on the vacuole for plant metabolism and development.

## INTRODUCTION

As sessile organisms, plants are facing frequent environmental fluctuations that constitute a challenge for their survival, growth and reproduction. Fluctuation in nutrient availability is one of the major factors limiting plant growth. Among those nutrients, nitrate is the major form of inorganic nitrogen taken up by plants in aerobic soil. As a critical nutrient for plant development, it is applied extensively in agriculture to sustain yields. However, because soil clay-humus complexes retain nitrate weakly, it is easily leached thereby leading to severe environmental pollution (Strahm and Harrison, 2006). One of the current challenges of plant breeding is therefore to generate crop varieties with imporved nitrogen use efficiency (NUE) to reduce excessive effluents in river and underground water.

Nitrate is absorbed by the roots and translocated to the shoot (Dechorgnat et al., 2011; Cookson et al., 2005). Once inside the cells, its assimilation occurs through the combined actions of different enzymes: nitrate reductase (NR) converts the nitrate intro nitrite that is reduced into ammonium by nitrite reductase (NiR). The synthetized ammonium is then incorporated into amino acids through the glutamine synthetase/glutamate synthase (GS/GOGAT) cycle. At the cellular level, plants are able to adjust their cytosolic nitrate concentrations between 1 and 6 mM according to nitrate availability in the environment (Miller and Smith, 2008; Cookson et al., 2005; Demes et al., 2020). The regulation of nitrate assimilation is essential to achieve such homeostasis. In parallel, the vacuolar compartment also plays a key role in the fine-tuning of cytosolic nitrate concentrations. When the external concentrations of nitrate are high, plants store it in their vacuole from which it can be remobilized when demand increases, as during a starvation period (Miller and Smith, 2008; Martinoia et al., 1981). To accumulate nitrate at high concentrations in the vacuole, the presence of an active transport is required (Miller and Smith, 1992). It was early suggested that this transport is mediated by an antiporter energized by vacuolar proton pumps generating pH gradient through the vacuolar membrane (Schumaker and Sze, 1987). Such a vacuole localized 2NO_3_^-^/H^+^ exchanger called AtCLCa was characterized by electrophysiological measurements on *Arabidopsis thaliana* isolated vacuoles (De Angeli et al., 2006). A knock- out for AtCLCa (*clca-2)* displays a decrease by up to 50% of the endogenous nitrate content, supporting its major role for nitrate storage in the vacuole (De Angeli et al., 2006).

Consequently, the reduction of AtCLCa activity in this mutant leads to an increase in nitrate assimilation and a change in root nitrate influx to adjust cytosolic nitrate homeostasis (Monachello et al., 2009; Liao et al., 2018).

It is assumed that nitrate in the vacuole does not only ensure nitrate homeostasis and proper plant growth under starvation but also plays a role as an osmoticum involved in plant water homeostasis (McIntyre, 1997). Genetic approaches support this hypothesis as QTLs for nitrate and water contents in non-limiting nitrogen conditions co-localize (Loudet et al., 2003). The *AtCLCa* gene is highly expressed in mesophyll cells and stomata. In the *clca-2* KO mutant, stomata opening in response to light and closure to abscisic acid (ABA) are impaired suggesting that AtCLCa is involved in anion translocation through the vacuolar membrane in both directions depending of the environmental conditions. Consequently the *clca-2* mutant is highly sensitive to hydric stress compared to wild-type plants (Wege et al., 2014), supporting a central function of AtCLCa in the control of water content regulation.

AtCLCa is a member of a highly conserved protein family widespread from prokaryotes to mammals (Mindell and Maduke, 2001). Although most CLCs are more selective for chloride, AtCLCa transports mainly nitrate as its selectivity motif contains a proline instead of the serine found in other characterized CLC exchangers (De Angeli et al., 2006; Wege et al., 2010). Additionally, despite their close structural similarity, CLC members can be either anion channels or anion/proton exchangers. In humans, five CLCs are chloride/proton exchangers (HsCLC3 to 7) whereas the four others are chloride channels (HsCLC1, HsCLC2, HsCLCΚa, HsCLCΚb) (Poroca et al., 2017). Interestingly, most of the exchangers share a highly conserved glutamate residue (E203 in AtCLCa), the “gating glutamate”. This residue, initially identified in CLC-ec1 from *E.coli* (Dutzler et al., 2002), is located in CLCs’ selectivity filter and projects its side chain in the ion pathway. When deprotonated, this residue blocks anion transport but, upon protonation, it moves out of the pathway thereby allowing anion access (Dutzler et al., 2003; Park et al., 2017). During a protonation/deprotonation cycle of this residue, two anions can be transported by the exchanger. The mutation of this glutamate in a non-protonable residue in bacteria (CLC-ec1), human (CLC-3, CLC-5 and CLC-7) (Costa et al., 2012; Weinert et al., 2010; Novarino et al., 2010; Weinert et al., 2020) and plant CLCs (AtCLCa) (Bergsdorf et al., 2009) uncouples anion transport from the proton transport and converts the exchanger into a channel.

In mammals, both CLC exchangers and channels coexist (Poroca et al., 2017). Interestingly, CLC transporters are all localized in intra-cellular compartments whereas, CLC channels are restricted to the plasma membrane. In Arabidopsis, all CLCs are localized in intracellular compartments and, so far, no CLC channel has been identified. In addition to AtCLCa, three other CLCs are located in the vacuolar membrane in *A. thaliana*. Among them, AtCLCb, the closest homologue of AtCLCa, is also a 2NO_3_ ^-^/H^+^ exchanger (Lv et al., 2009; von der Fecht-Bartenbach et al., 2010). Nevertheless, knock-out mutants for *AtCLCb* contain as much nitrate as the wild-type genotype suggesting that loss of AtCLCb is compensated by AtCLCa (von der Fecht-Bartenbach et al., 2010). The other vacuolar CLCs in Arabidopsis, AtCLCc and AtCLCg, are involved in chloride transport, as the knock-out mutants are more sensitive to NaCl stress, but their electrophysiological properties are unknown to date (Jossier et al., 2010; Nguyen et al., 2016).

AtCLCa is thus an essential transporter for nitrate storage in the vacuole and the control of water content. As an exchanger mechanism was demonstrated for AtCLCa, we wondered if nitrate and proton transport coupling is absolutely required for plants to stabilize water and nitrate status. We investigated this question by analyzing the physiological consequences of a conversion of the AtCLCa exchanger into a channel. We mutated the gating glutamate of AtCLCa into an alanine, a non-protonable residue (E203A), and introduced it in a knock-out *clca* background to analyze the phenotype of the generated plants for water content and nitrogen use efficiency. The physiological consequences of such a mutation should provide insight on the significance of having an exchanger rather than a channel activity for AtCLCa.

## RESULTS

### Expression of AtCLCa with a gating glutamate mutation in *clca* KO plants

To analyze the physiological consequences of the E203A mutation in AtCLCa, *AtCLCa_E203A_* under the control of the 35S promoter or the AtCLCa native promoter was introduced in *clca- 2* knockout mutant (De Angeli et al., 2006). As a positive control, we used the complemented line, *clca-2/35S:AtCLCa*, already characterized in previous studies (Wege et al., 2010, 2014) and two control lines *clca-2/pAtCLCa:AtCLCa* generated in this study. The *clca- 2/35S:AtCLCa_E203A_ 3* and *8* lines were selected because they overexpress AtCLCa as strongly as *clca-2/35S:AtCLCa* complemented plants (20 to 40 fold relative to WT), whereas *clca- 2/pAtCLCa:AtCLCa_E203A_ 1* and *4* lines and the *clca-2/pAtCLCa:AtCLCa 6* and *2* lines were selected as they display an expression level 0.5 to 2 fold compared to native *CLCa* in Ws-2 (Supplemental Figure S1A). In parallel, we checked that the mutation in AtCLCa does not change the sub-cellular localization of the protein by transforming *clca-2* mutant with *clca- 2/35S:GFP-AtCLCa_E203A_*. The fluorescence was observed on plants from two different independent lines in guard cells and apical root cells (Supplemental Figure S1B). As expected, the mutated form of AtCLCa is localized in the vacuolar membrane in both cell types.

### AtCLCa_E203A_ shows reduced proton/anion coupling

A previous report showed that in Xenopus oocytes the “gating glutamate” mutation E203A in AtCLCa exchanger disrupts NO_3_^-^/H^+^ coupling (Bergsdorf et al., 2009). Therefore, to quantify the changes in the vacuolar anion transport induced by the E203A mutation in AtCLCa, we investigated the properties of the ion currents across mesophyll vacuolar membranes from the *clca-2/35S:AtCLCa_E203A_* L3 and L8 lines (Figure 1). We applied the patch-clamp technique to vacuoles from these two genotypes as well as *clca-2/35S:AtCLCa,* Ws-2 and *clca-2* knock out mutant in the whole-vacuole configuration. In order to measure anionic currents only, we used the non-permeable cation BisTrisPropane as a counter ion. We found that *clca- 2/35S:AtCLCa_E203A_* L3 and L8 lines had similar behavior (Figure 1A compared to Supplemental Figure S2A), thus for detailed characterization we focused on *clca- 2/35S:AtCLCa_E203A_ L3* line. In order to evaluate the impact of the E203A mutation on the H^+^/NO_3_^-^ coupling of AtCLCa and the intensity of NO_3_^-^ currents across the tonoplast of the different transgenic lines, we used the experimental design schematized in Figure 1A. First, we exposed vacuoles to bi-ionic condition (i.e. NO_3_^-^ in the vacuole, Cl^-^ on the cytosolic side) to measure the vacuolar current densities in the different genotypes. Second, vacuoles were exposed to NO_3_^-^ in the cytosol (i.e. with NO_3_^-^ on both sides of the vacuolar membrane) to allow comparison between the Nernst equilibrium potential for NO_3_^-^ (E_Nernst_^NO3^) and the measured reversal potential (E_rev_) (De Angeli et al., 2006). Third, to test the coupling between NO_3_^-^ and H^+^ transports, the cytosolic pH was shifted from 7 to 9 in presence of NO_3_^-^ at the cytosolic side to quantify the change in E_rev_. Finally, each vacuole was exposed to the initial bi-ionic conditions to ensure that it was not damaged by the treatments.

**Figure 1.**
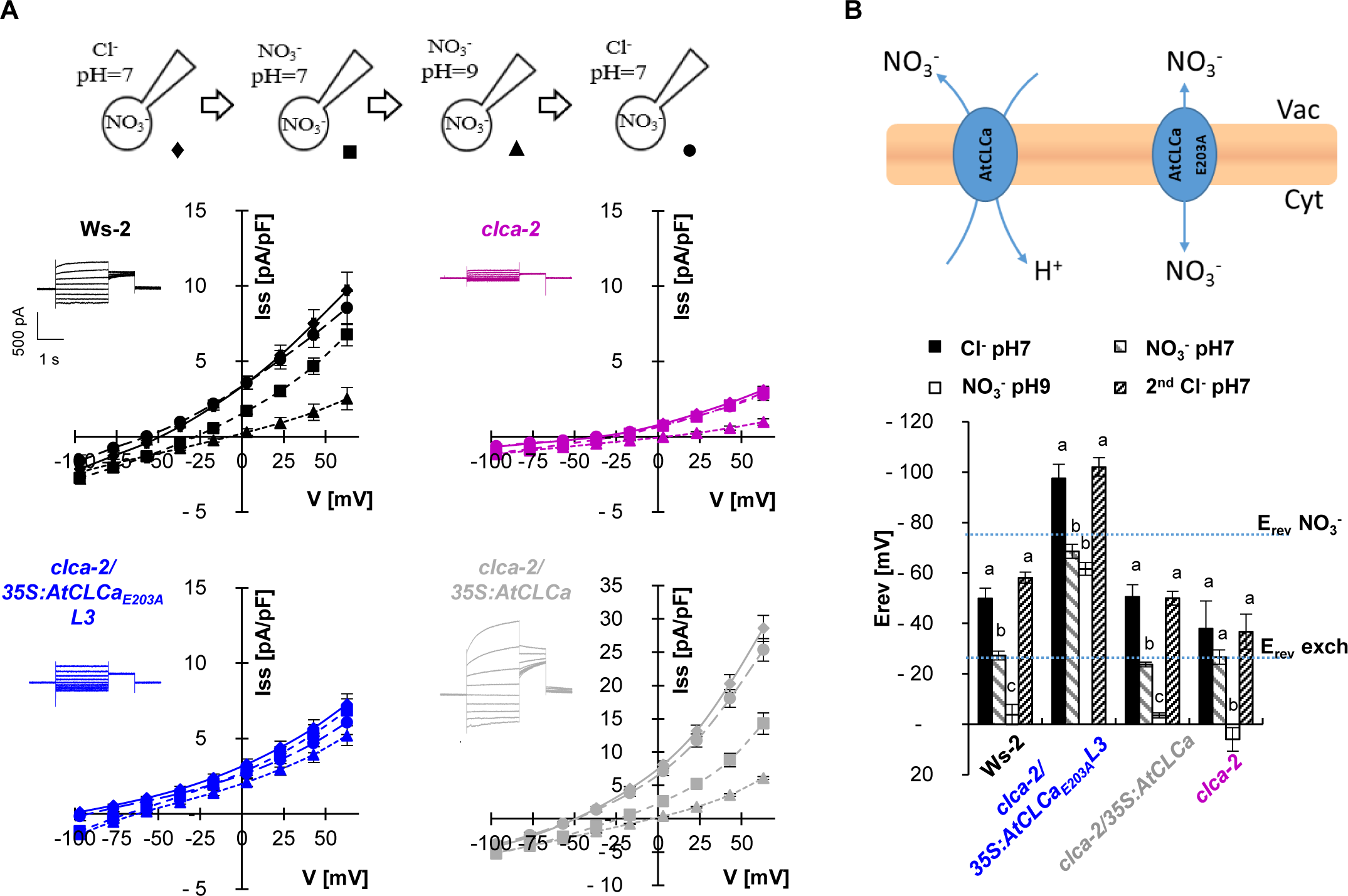
*AtCLCa_E203A_* overexpression restores tonoplast anion currents but alters the pH dependency. (A) Steady state current density (Iss) from Ws-2, *clca-2, clca-2/35S:AtCLCa_E203A_ L3* and *clca-2/35S:AtCLCa* vacuoles in standard conditions containing 20mM Cl^-^ pH7 (♦/ return ●), 4.2mM NO_3_ ^-^ pH7 (▪) and 4.2mM NO_3_ ^-^ pH9 (▲) in the extra vacuolar media were measured (A). Iss were plotted against the applied membrane potential. Representative whole-vacuole currents from each genotype in standard conditions are displayed in small figures. (B) The reversal potentials (Erev) of the four genotypes were recorded in all measured conditions and reveal elevated Erev in *clca-2/35S:AtCLCa_E203A_ L3* vacuoles, close to Erev for a nitrate channel when the one from Ws-2 confirms that WT AtCLCa is a NO_3_ ^-^/H^+^ exchanger. Only stable measurements of vacuoles that returned to initial reversal potentials in the starting conditions (2^nd^ Cl^-^ pH7) were considered. Data represents means ±SEM of n≥5 vacuoles of at least 4 different plants. One-way ANOVA analysis with Bonferroni comparison post-test (p<0.05) were applied, different letters indicate significant difference inside each genotype.

As previously shown, in vacuoles from *clca-2* the current density was much lower than in the wild type Ws-2 or in *clca-2/35S:AtCLCa*, corresponding to a decrease by 66 ± 7 % and 89 ± 1 % at +43 mV under bi-ionic conditions (20 mM Cl^-^ pH 7 at the cytosolic side), respectively (Figure 1A). E_rev_ in *clca-2* was difficult to quantify due to high variance (Supplemental Figure S2B) probably resulting from the very low vacuolar current densities measured in this genotype. Under all tested conditions, the currents mediated by AtCLCa_E203A_ were twice higher than in *clca-2*. Notably, in AtCLCa_E203A_ vacuoles, no activating kinetics of the ion currents at positive membrane potentials could be observed (Figure 1A), suggesting a link between activation at positive membrane potential and the exchanger mechanism of AtCLCa. Plotting the measured steady-state current densities (Iss) against the applied voltage revealed, in all ionic conditions, a far more negative reversal potential (E_rev_) for *clca- 2/35S:AtCLCa_E203A_* vacuoles compared to Ws-2, *clca-2/35S:AtCLCa* and *clca-2* vacuoles (Figure 1). While the E_rev_ of Ws-2 and *35S:AtCLCa* measured when nitrate is in the cytosol confirmed the previously reported 2NO_3_^-^/1H^+^ transport stoichiometry (De Angeli et al., 2006), in *clca-2/35S:AtCLCa_E203A_* vacuoles we observed a reversal potential of -68.5 ± 6.3 mV that is close to the E_Nernst_^NO3^ = -75 mV. The proximity of the E_rev_^E203A^ with E_Nernst_^NO3^ indicates that the coupling of anion and H^+^-transport is dramatically affected in AtCLCa_E203A_. In the next step, the change of pH from 7 to 9 at the cytosolic side of the vacuolar membrane confirmed the disruption of the H^+^ coupling in AtCLCa_E203A_. The cytosolic pH changes significantly modified the measured reversal potentials in Ws-2 (E_rev_^pH7^= -27.2 ± 4. mV and E_rev_^pH9^= -3.7 ± 10.0) and in *clca-2/35S:AtCLCa* (E_rev_^pH7^= -23.7 ± 2.0 mV and E_rev_^pH9^= -3.4 ± 2.2 mV). Notably, the ΔE_rev_ observed in Ws-2 (ΔE_rev_= +23.5 ± 6.7 mV) and *clca- 2/35S:AtCLCa* (Δ E_rev_ = +20.2 ± 2.7 mV) is close to the expected shift for a 1H^+^/2NO_3_^-^ antiporter. In contrast, in AtCLCa_E203A_ vacuoles the E_rev_ after exposure to cytosolic side pH 9 was not significantly affected (E_rev_^pH7^= -68.5 ± 6.3 mV and E_rev_^pH9^= -61.6 ± 5.7 mV) confirming the absence of H^+^ coupling in AtCLCa_E203A_ mutants (Figure 1B). These data demonstrate that the expression of *35S:AtCLCa_E203A_*in *clca-2* mutant does not restore 1H^+^/2NO_3_^-^ antiporter activity in the vacuolar membrane. Further, in *clca-2/35S:AtCLCa_E203A_*we observed a higher current density compared to *clca-2.* Therefore, from this set of data we can conclude that the *clca-2/35S:AtCLCa_E203A_* plants express a passive NO_3_^-^ selective transport system in the vacuolar membrane that is absent in *clca-2* and distinct from the 1H^+^/2NO_3_^-^ antiporter activity detected in the other genotypes.

### Expression of AtCLCaE203A in *clca* mutant does not restore plant growth

Nitrate has been known for decades to be a crucial nutrient for plant growth, notably because of its involvement in nitrogen metabolism (Brouwer, 1962; Crawford, 1995; Chen et al., 2004). We therefore analyzed the consequences of the introduction of the mutation AtCLCa_E203A_ on plant growth. After 6 weeks of growth on 4.25 mM NO_3_ ^-^ in short day conditions, the fresh weight of *clca-2* mutant shoot was decreased by 30 ± 5 % compared to Ws-2 plants (Figure 2B). The introduction of *AtCLCa* restored the wild-type phenotype irrespective of whether the endogenous or the 35S promoter was used to drive its expression (Figure 2 and Supplemental Figure S3A). Surprisingly, not only expression of *AtCLCa_E203A_*did not rescue *clca-2* phenotype, but also *clca-2/35S: AtCLCa_E203A_*plants shoot and root fresh weights were even further decreased by 26 ± 5 % and 29 ± 7 % compared to *clca-2* (Figure 2B). The shoot to root fresh weight ratio was not affected in any of the phenotypes, indicating that the plants were not nutrient-starved (Castaings et al., 2009; Lawlor et al., 2001). Under the control of the endogenous promoter, *AtCLCa_E203A_* expression in *clca-2* did not rescue plant shoot fresh weight either (Supplemental Figure S3A). However, only one of the two *clca- 2/pAtCLCa:*AtCLCa_E203A_ lines analyzed displayed a statistically significant plant fresh weight reduction (22 ± 4 %) compared to *clca-2*. In conclusion, *AtCLCa_E203A_* expression is not able to rescue the growth deficiency phenotype of *clca-2* in the tested conditions and even exacerbates it when over-expressed ubiquitously.

**Figure 2.**
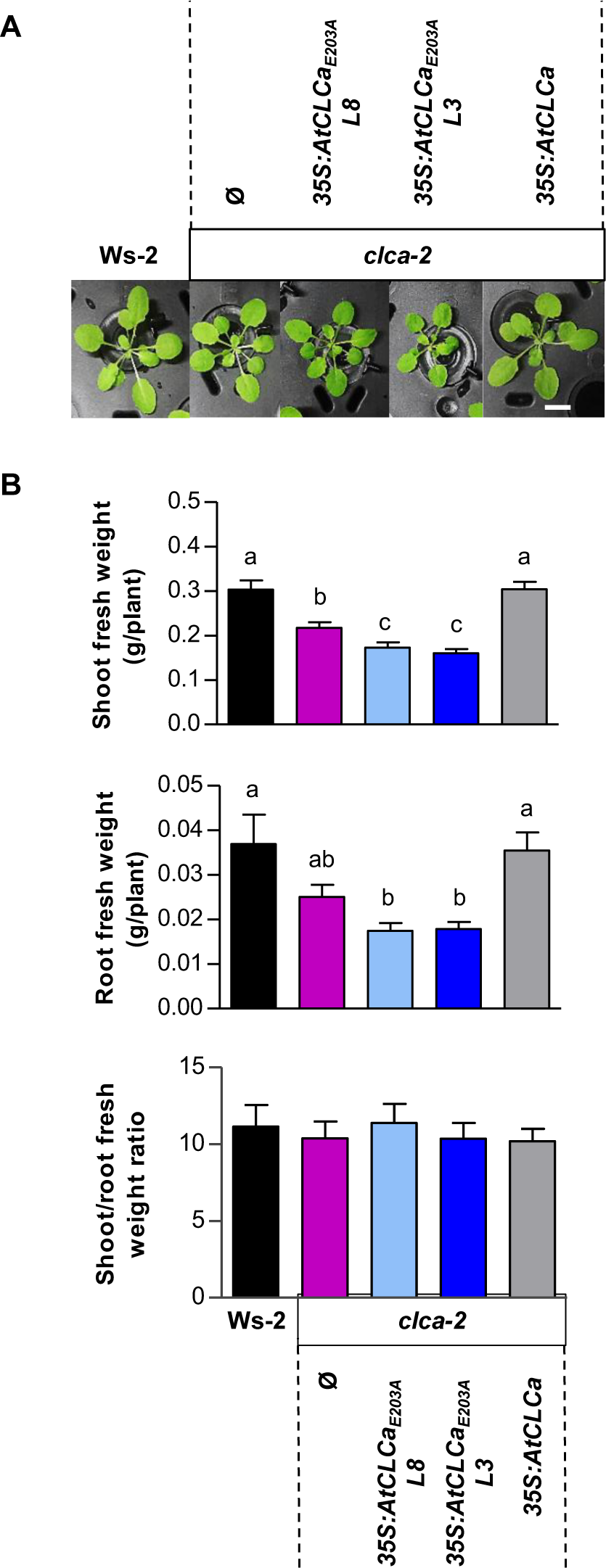
*AtCLCa_E203A_* does not complement *clca-2* biomass production deficiency. Plants of Ws-2, *clca-2*, *clca-2/35S:AtCLCa_E203A_* lines 8 and 3 and complemented line *clca-2/35S:AtCLCa* were grown in hydroponics on 4.25 mM NO_3_ ^-^ under short day conditions. After four weeks, photographs were taken **(A)** and after six weeks shoot and root fresh weights and the shoot/root ratio biomasses were measured **(B)**. Data represent the means ± SEM of three biological replicates (3<n<6 per replicate). A Shapiro-Wilk normality test followed by a Welch’s t-test were applied. Different letters indicate significant difference between genotypes (p<0.05). Scale bar represents 1 cm.

### Water homeostasis is disrupted in plants expressing *AtCLCa_E203A_*

In order to understand why the E203A form of AtCLCa leads to a decrease in plant growth when over-expressed, we explored the impact of uncoupling AtCLCa on plant water homeostasis. Indeed, nitrate is not only an essential nutrient but also a major signaling molecule and an important osmoticum for plant cells (McIntyre, 1997; Wege et al., 2014). *AtCLCa* is expressed in both mesophyll and guard cells where it is involved in building up the osmotic potential required for proper stomata movements (Wege et al., 2014). We first measured stomata opening in response to light in plants over-expressing *AtCLCa_E203A_*(Figure 3A). As shown previously, stomata opening is impaired in *clca-2* (Wege et al., 2014). In plants overexpressing *AtCLCa_E203A_*, interestingly, we observed two phases: first, the opening followed the same kinetics as in *clca-2* and, after 120 minutes, in a second phase, the stomata opening became significantly lower in *clca-2/35S:AtCLCa_E203A_* compared to *clca-2.* Similar results were obtained for the lines with the constructs *pCLCa:AtCLCa_E203A_* (Supplemental Figure S4A). Therefore, AtCLCa exchanger mechanism is required for efficient stomata opening in response to light. We also investigated stomata closure induced by ABA on epidermis peels. As expected, stomata from *clca-2* responded very weakly to this hormone (Wege et al., 2014). In *clca-2/35S:AtCLCa_E203A_*plants, ABA-induced stomata closure was reduced to a similar extent as in *clca-2* (Figure 3B, Supplemental Figure S4A). Thus, these results show that the uncoupled AtCLCa responds very weakly to ABA and affects both stomata opening and closure. This indicates that the exchanger mechanism is essential for the control of the ionic and, consequently, water fluxes through the vacuolar membrane, which are necessary for correct functioning of stomata.

**Figure 3.**
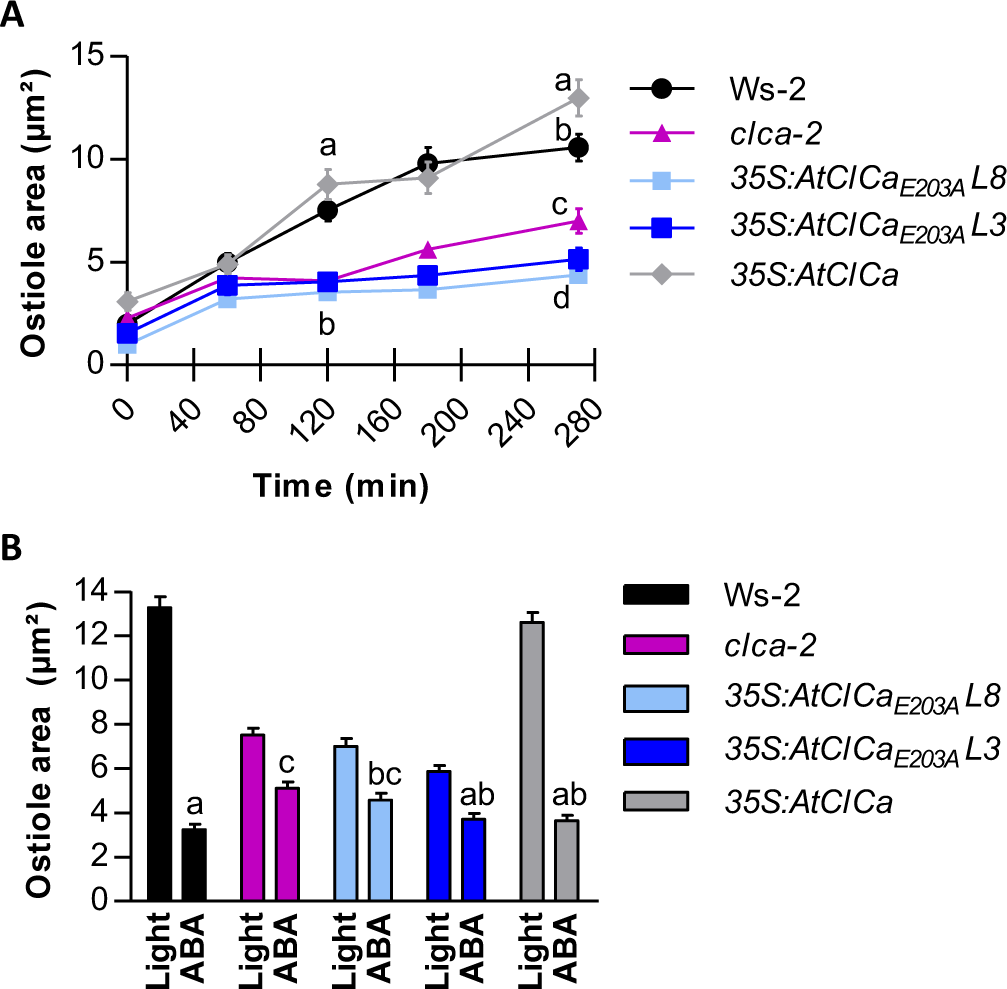
*AtClCa_E203A_* plants are affected in stomata movements. Kinetics of stomata opening in response to light **(A)** and effect of ABA on stomata closure **(B)**. Experiments were performed on isolated epidermal peels of five weeks plants grown as in Figure 1. Epidermis were incubated in KCl buffer for 1 hour in the dark before their transfer in light for 4.5h followed by a 50 µM ABA treatment for 3h. Data represent the means ± SEM of three biological replicates (n=85-150 per replicate). One-way ANOVA analysis with Bonferroni comparison post-test (p<0.05), different letters indicate significant difference.

To further analyze the consequences of the mutation in AtCLCa on water content in whole Arabidopsis, plants were grown on well-watered soil under short day conditions. In *clca-2* mutant, the dry weight and water content were significantly lower compared to Ws-2 (Figure 4A). This phenotype was restored by overexpressing AtCLCa as previously shown (Wege et al., 2014). However, overexpression of *AtCLCa_E203A_* did not allow rescuing wild-type dry weight and water content. It even led to a further reduction of the water content compared to *clca-2* indicating that the coupling of nitrate and proton transport in AtCLCa is required for water homeostasis. Expression of *AtCLCa_E203A_* under the control of its endogenous promoter also failed to rescue the wild-type water content (Supplemental Figure S3B). However, only one line displayed a further decrease of water content compared to *clca-2* thereby indicating that the overexpression is responsible for the aggravation of the *clca-2* growth phenotype in *clca-2/35S:AtCLCa_E203A_*.

**Figure 4.**
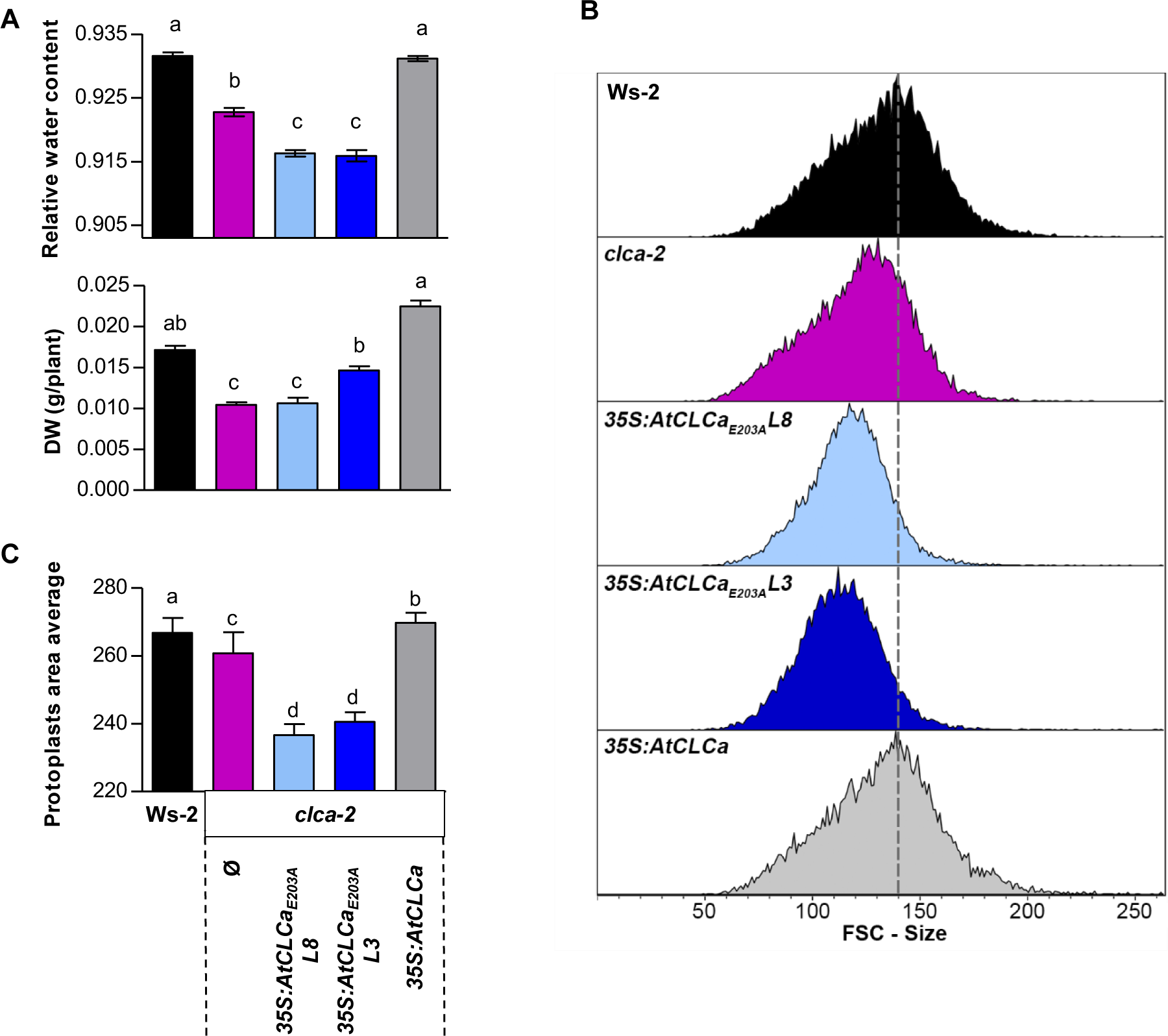
*AtCLCa_E203A_* plants contain less water leading to impaired cell enlargement. **(A)** Relative water content of six weeks plants, grown for five weeks on soil under short day conditions, expressing *AtCLCa_E203A_* under the control of the 35S promoter. Three biological replicates (25<n<30 by replicate). Statistical analysis as in Figure 3. **(B)** Distribution of relative cell sizes determined by FACS (flow cytometry) of protoplasts generated from leaves enzymatic digestion of five weeks old plants grown as in A. The data presented are representative one experiment. **(C)** Mean of the relative cell sizes obtained by FACS for each genotype. Two biological replicates (n=3 plants per replicate, protoplasts > 30 000). Statistical analysis are as in Figure 3.

As the importance of water for plant cell growth is well establised (Boyer, 1968), we decided to investigate the effect of expressing the uncoupled version of AtCLCa on cell size. This latter parameter was determined by flow cytometry on protoplasts produced by enzymatic digestion of leaves from plants over-expressing *AtCLCa_E203A_*. Chlorophyll detection allowed us to analyze specifically mesophyll cells and measure the distribution of the relative cell sizes for each genotype (Figures 4B and 4C). The size of mesophyll cells from both *clca-2* knock-out mutant was clearly reduced compared to Ws-2. Wild-type cell size was recovered upon expression of native AtCLCa. In contrast, cells from the *AtCLCa_E203A_*plants were even smaller than those from *clca-2* by up to 9.8 ± 3.7 % (Figures 4C). The decreased water content observed in plants affected in the gating glutamate of AtCLCa is correlated to a decrease of relative cell size, which could account for the lower fresh weight of those plants.

### Uncoupling NO_3_^-^ and H^+^ transport in AtCLCa modifies nitrate storage and remobilization kinetics

As AtCLCa was previously characterized for its function in nitrate storage (De Angeli et al., 2006), we analyzed the nitrate accumulation in plants expressing *AtCLCa_E203A_* under the control of the 35S promoter or the endogenous promoter. In agreement with previous works (Geelen et al., 2000; Monachello et al., 2009), we observed a decrease of 37 ± 4 % and 30 ± 4 % of the nitrate content in *clca-2* shoot and root respectively compared to Ws-2 (Figure 5A and Supplemental Figures S3C). Strikingly, *AtCLCa_E203A_* overexpression in *clca-2* induced a further decrease in nitrate content in shoots by 52 ± 4 % compared to *clca-2*, corresponding to a decrease by 70 ± 3 % compared to Ws-2. The nitrate content of *clca-2/35S:AtCLCa_E203A_*root was similar to *clca-2*. Expression of *AtCLCa_E203A_* under the control of its own promoter did not lead to further reduction of nitrate content compared to *clca-2* (Supplemental Figure S3C). As AtCLCa can still transport chloride but with a lower selectivity compared to nitrate (De Angeli et al., 2006), the content of this anion was measured (Figure 5B). In parallel, we analyzed the concentrations of other major anions and potassium that are not transported by AtCLCa but could be affected by the nitrate under-accumulation (Figures 5B to 5D). We found similar levels of these ions in the different genotypes. Nevertheless, *clca-2* displayed an increase of malate content as previously shown (Geelen et al., 2000). This increase was enhanced in *clca-2/35S:AtCLCa_E203A_*lines. Altogether, these results show that *AtCLCa_E203A_* is not able to restore wild-type phenotype for nitrate contents and indicate that uncoupling nitrate and proton transport in AtCLCa strongly alters nitrate storage into vacuoles.

**Figure 5.**
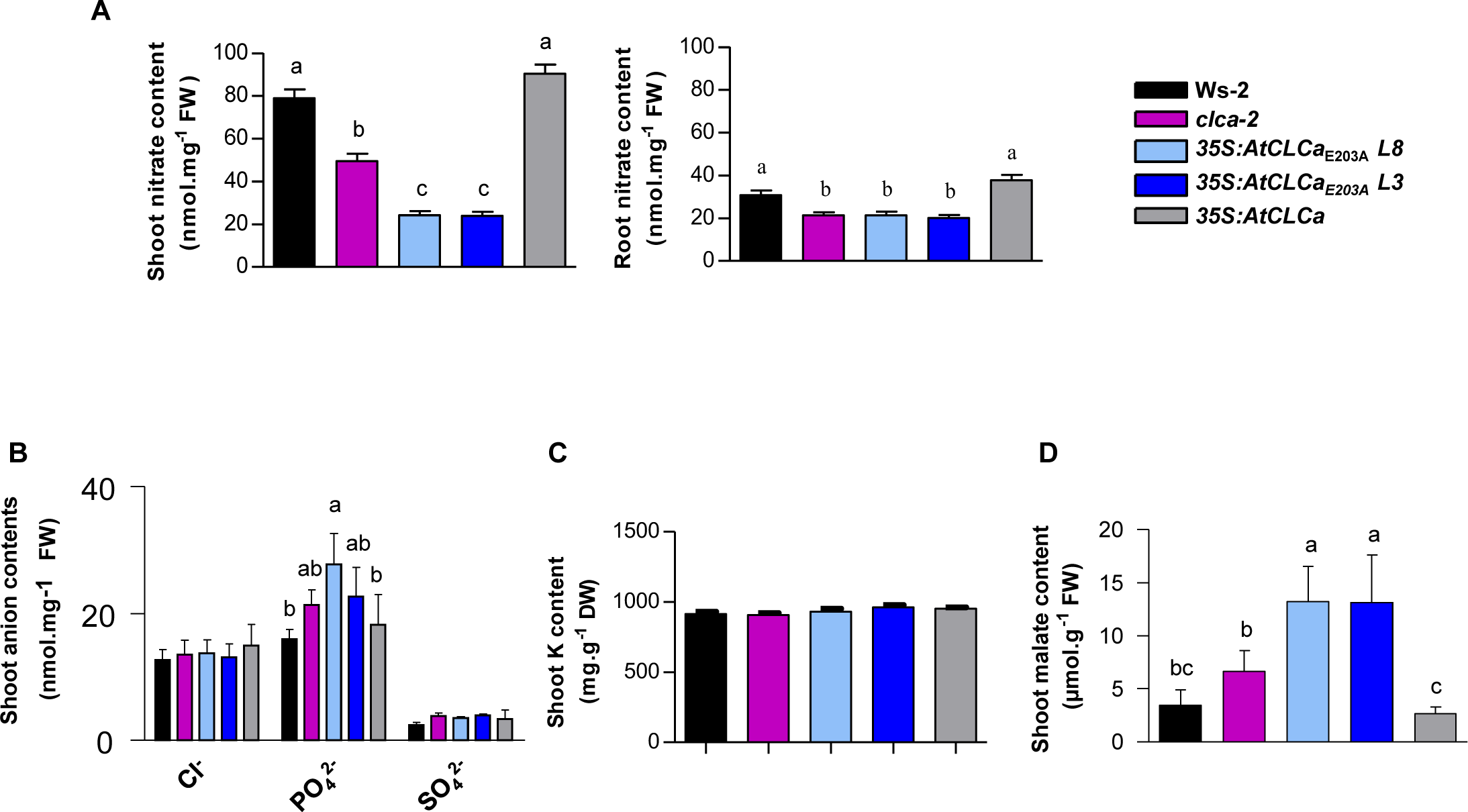
*AtCLCa_E203A_* expression indicates a decrease of endogenous nitrate but an increase of malate concentration. Endogenous nitrate contents **(A)** in shoot (left) and root (right), inorganic anion **(B)**, potassium **(C)** and malate **(D)** concentrations of Ws-2, *clca-2*, *clca-2*/*35S:AtCLCa_E203A_* and *clca-2/35S:AtCLCa* plant shoots grown in hydroponics as described in legend of Figure 2. Data represent the means ± SD of two experiments (with 2<n<9 by experiment). Statistical analysis as in Figure 3.

To better understand the phenotypes of the transgenic lines, we measured the kinetics nitrate accumulation and remobilization in response to nitrate availability which mainly reflects mainly the fluxes through the vacuolar membrane (Miller and Smith, 2008; Huang et al., 2012). We decided to focus for the following experiments only on plants overexpressing *AtCLCa_E203A_* in which nitrate content is particularly low compared to Ws-2 and *clca-2* whereas, the vacuolar anion currents driven by AtCLCa are nearly as high as in Ws-2. First, the dynamics of nitrate storage in plants expressing AtCLCa_E203A_ subjected to nitrate depletion was analyzed. Plants were grown for 5 weeks in hydroponics on complete Hoagland medium (4.25 mM NO_3_ ^-^) and then exposed to nitrogen starvation for 120 hours. The differences in nitrate content in the various lines at the beginning of the experiment corresponded to the ones observed in our previous tests (Figures 5A and 6A). In all genotypes, the kinetics of remobilization was similar during the first 72 hours of starvation either in the roots and the aerial parts, apart from the root of Ws-2: all plants lost around 0.25 nmol of nitrate per mg of fresh weight per hour (Figure 6A). This led to a complete nitrate depletion in roots of *clca-2* and *clca-2/35S:AtCLCa_E203A_* lines. Between 72 and 120 hours, the rates of nitrate remobilization in shoots increased leading to complete depletion of nitrate in *clca-2/35S:AtCLCa_E203A_* lines. As a control, we measured in parallel the chloride contents in the same plants (Supplemental Figure S5A). Nitrate starvation led to an increase of Cl^-^ content in the different genotypes and an over-accumulation in roots of *clca- 2/35S:AtCLCa_E203A_* compared to Ws-2 and *clca-2*. Then this experiment indicated that: (1) the plants adjust the deficiency of negative charges linked to the absence of nitrate by stimulating the absorption of chloride and, (2) the net rate of nitrate remobilization is not affected by the lack of AtCLCa or the presence of AtCLCa_E203A_ on the vacuolar membrane. The time to reach complete depletion is essentially related to the level of nitrate stored at the beginning of the experiment in the different genotypes and organs.

**Figure 6.**
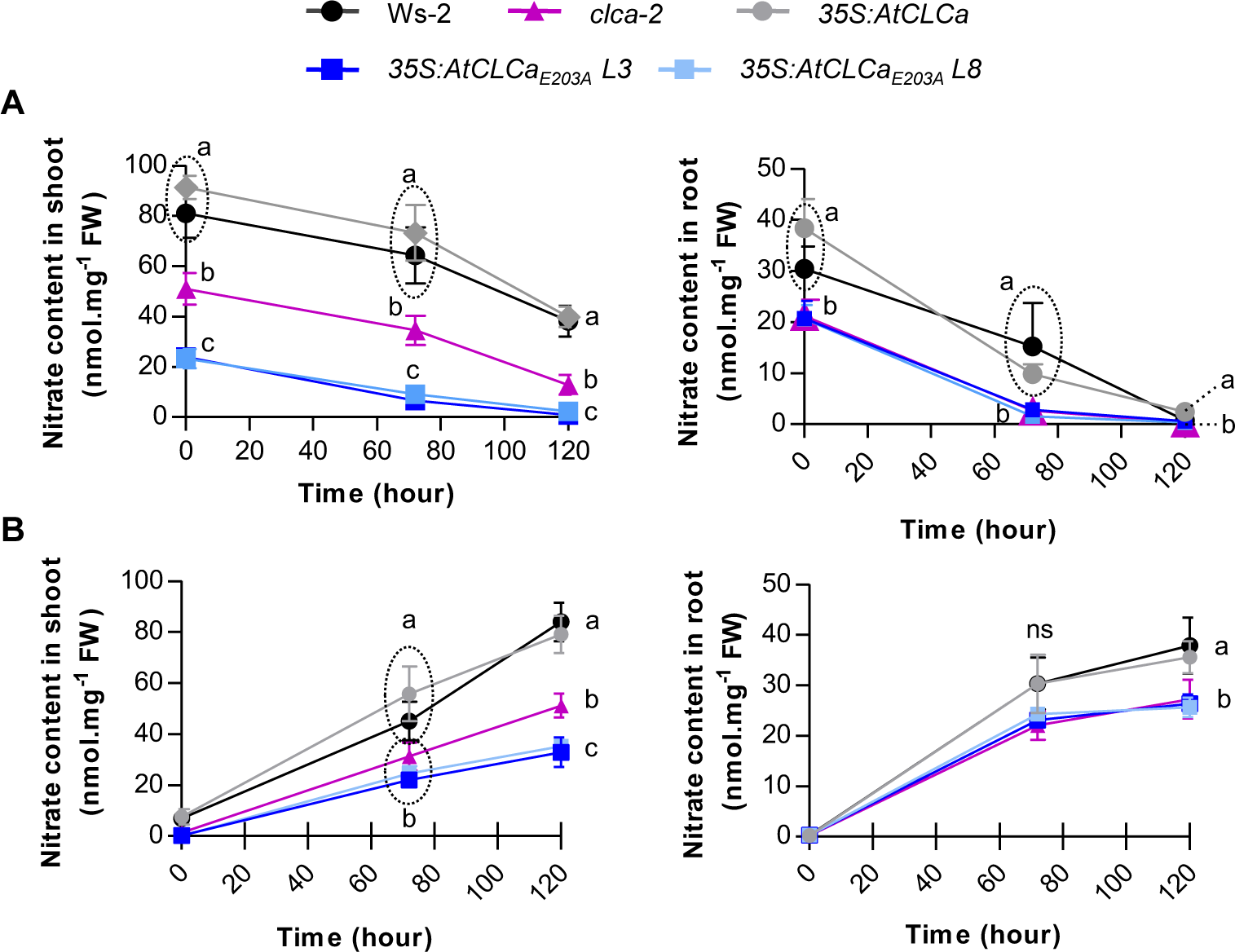
*clca-2/35S:AtCLCa_E203A_* plants present a slower nitrate storage and no change in nitrate remobilization in response to nitrate supply in the medium compared to wild-type. **(A)** Plants were cultivated in hydroponics for five weeks as described in Figure 2 and then nitrogen was removed from the medium for 120h. Nitrate content was determined after 0, 72h and 120h of starvation. **(B)** Five week-old plants were submitted to ten days of nitrogen starvation, and 4.25 mM nitrate was supplied again, the content was determined after 0, 72h and 120h. Nitrate concentration is quantified in shoots (left) and in roots (right) separately. In both experiments, data represent the means ± SEM of three biological experiments (n=4-6 plants per replicate). Statistical analysis as in Figure 3. ns: not significant.

To investigate the kinetics of nitrate accumulation in the vacuole, plants were nitrogen starved for ten days leading to nitrate concentrations close to zero in all genotypes, afterwards 4.25 mM nitrate was resupplied to the plants and nitrate accumulation was measured (Figure 6B). In all genotypes, nitrate content increased when this anion was added to the medium whereas chloride concentrations decreased showing again a negative correlation between the quantities of these two anions *in planta* (Supplemental Figure S5B). Nevertheless, chloride slightly over-accumulated after 120 hours only in shoots by 23.5±1.0 % in *clca-2/35S:AtCLCa_E203A_*lines compared to Ws-2 and *clca-2*. In shoots, at the end of the kinetics, a significant difference in the rate of nitrate accumulation between *clca-2* and *clca-2/35S:AtCLCa_E203A_*lines was obvious: rates of 0.65 nmol NO_3_^-^ mg FW^-1^ h^-1^ were measured for Ws-2 and *clca- 2/35S:AtCLCa* whereas they were only 0.4 and 0.25 NO_3_^-^ mg FW^-1^ h^-1^ were observed for *clca-2* and *clca-2/35S:AtCLCa_E203A_* lines. These differences led to lower nitrate accumulation of 39.1±9.7 % and 60.5±8.1 % compared to wild-type in *clca-2* and the two transgenic lines, respectively. In roots, the storage rate decreased by up to 24±4 % compared to Ws-2 but was similar to the one measured in *clca-2*. These results indicate that, the introduction of the uncoupled form of AtCLCa in *clca-2* strongly decreases the rate of nitrate accumulation into the vacuole explaining the difference of nitrate contents in the different lines.

### AtCLCa_E203A_ enhances nitrate assimilation and nitrogen use efficiency

The defect in nitrate storage into the vacuole in the plants expressing AtCLCa_E203A_ is likely to unbalance the cytosolic nitrate homeostasis and to consequently change nitrate assimilation. To test this hypothesis, we measured the activity of nitrate reductase (NR), the first enzyme involved in nitrate assimilation localized in the cytosol, on four week-old *clca- 2/35S:AtCLCa_E203A_* plants in which nitrate storage is the most reduced (Figure 7A). The analysis was performed three hours after the dark-light transition when the activity is the highest (Man et al., 1999). The NR activity increased by 4-fold in *clca-2* and 6.5-fold in *clca- 2/35S:AtCLCa_E203A_* compared to wild-type. Consequently, the total amount of free amino acids was higher by up to 42 ± 2 % in the 2 transgenic lines overexpressing *AtCLCa_E203A_*(Figure 7B). However, the total amount of free amino acids was not different between Ws-2 and *clca-2* KO mutant. Interestingly, asparagine (Asn), serine (Ser), glutamine (Gln) and glycine (Gly) were significantly accumulated in *AtCLCa_E203A_*lines while the concentrations of other amino acids did not change, indicating a modification of amino acid distribution (Figure 7B and Supplemental Figure S6). We wondered whether the increase in free amino acid concentration induced by uncoupling nitrate and proton transport in AtCLCa affects the protein content is also affected. No significant difference was observed between the wild-type, *clca-2* and *clca-2/35S:AtCLCa*. Interestingly, *clca-2/35S:AtCLCa_E203A_* plants displayed an increase in protein content (25 ± 5 %) compared to Ws-2 (Figure 7C). Those results suggest that inefficient vacuolar nitrate storage due to *AtCLCa_E203A_* overexpression leads to increased nitrate assimilation into amino acids and proteins.

**Figure 7.**
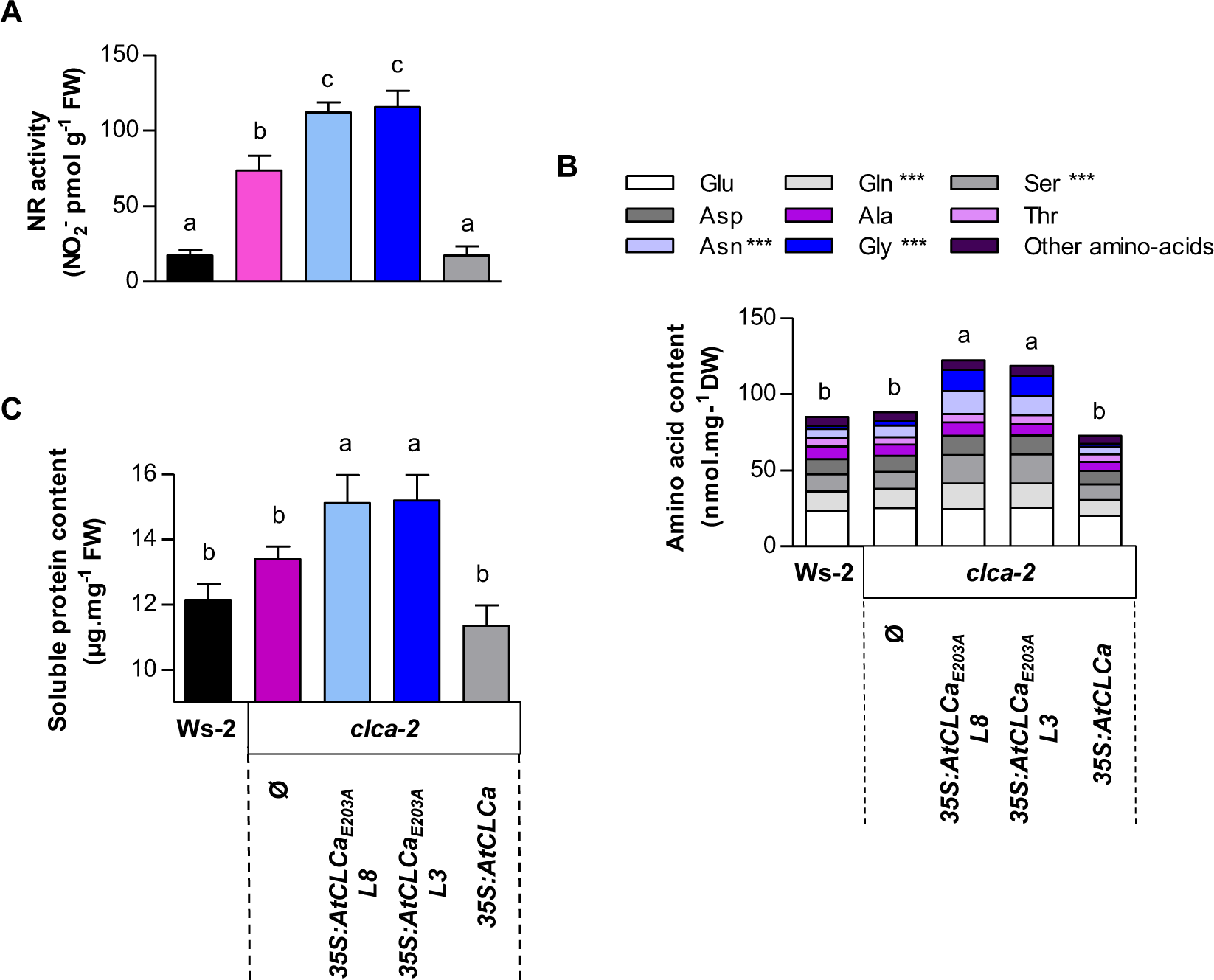
*clca-2/35S:AtCLCa_E203A_* plants have an increased nitrate assimilation at vegetative stage. Plants of Ws-2, *clca-2*, *clca-2/35S:AtCLCa_E203A_ line 8* and *3* and *clca-2/35S:AtCLCa* were grown as described in Fig 2 and analyzed for their nitrate reductase (NR) activity **(A)**, total free amino-acids and individual amino-acids contents **(B)** and soluble protein content **(C)**. For nitrate reductase activity, the analysis was performed on four to five plants after 3 hours to light. For amino-acids content, four plants were analysed, stars represent significant differences of absolute amino acid content between *clca- 2/35S:AtCLCa_E203A_* and Ws-2, *clca-2* and *clca-2/35S:AtCLCa.* For soluble protein content, three biological replicates including three plants were performed. Data represent the means ± SD, statistical analysis is the same as in Figure 3.

Based on these findings, we wondered whether nitrogen metabolism is also perturbed by AtCLCa_E203A_ overexpression at later developmental stages. To this aim, we determined the nitrogen use efficiency (NUE) in the five genotypes used above. The plants were labelled with ^15^N at the grain filling stage. At harvest, no difference in total dry weight between the genotypes was observed except for the transgenic line *clca-2/35S:CLCa_E203A_* L3. Nevertheless, dry weight partitioning between rosettes, stems and roots was the same in all genotypes (Supplemental Figure S7). This result confirms that the alteration of fresh weight noticed previously between wild-type and transgenic plants at the vegetative stage is mainly due to a change in water status (Figure 4). Nitrogen content analysis allowed us to quantify N allocation in the aboveground organs. Knock-out *clca-2* mutant retained 18.3 % less N than Ws-2 in rosettes while its seeds were enriched in N by 5.5 % (Figure 8A). N partition in rosette leaves was even lower in the two *clca-2/35S:CLCa_E203A_*lines (29.5% and 13.7% compared to Ws-2 and *clca-2* respectively), which resulted in much higher N partition in seeds compared to Ws-2 and *clca-2* leading to a higher N concentration in seeds (Figure 8B). This increase in nitrogen concentration in seeds cannot be explained by a difference in N remobilization efficiency (NRE) between the source and sink organs. Indeed, the values of this parameter, corresponding to the partitioning of ^15^N in seeds at harvest compared to the harvest index (^15^NHI/HI; Chardon et al., 2012), were not different among the five genotypes (Figure 8C). In parallel, we estimated the relative specific absorption ratio (RSA ratio), corresponding to the ratio between ^15^N in seeds at harvest and the N harvest index (^15^NHI/NHI), indicating the dilution of ^15^N in seeds due to the post-flowering uptake (Chardon et al., 2012). The values showed a decrease between Ws-2 and *clca-2* or *clca- 2/35S:AtCLCa_E203A_* (Figure 8D). These differences may account for differences in N allocation through the different aerial organs of the plants observed previously. Finally, we estimated NUE for grain production, calculating the ratio between the proportion of nitrogen allocated to seeds and the harvest index (NHI/HI; Marmagne et al., 2020). In Ws-2, the value of NUE was 1.21 ± 0.18. In *clca-2*, it increased to 1.30 ± 0.14 and reached 1.43 ± 0.22 in *clca-2/35S:CLCa_E203A_* plants (Figure 8D). These results indicate that mutating the gating glutamate in AtCLCa leads to a higher allocation of N uptake to seeds and consequently to a better NUE at reproductive stage.

**Figure 8.**
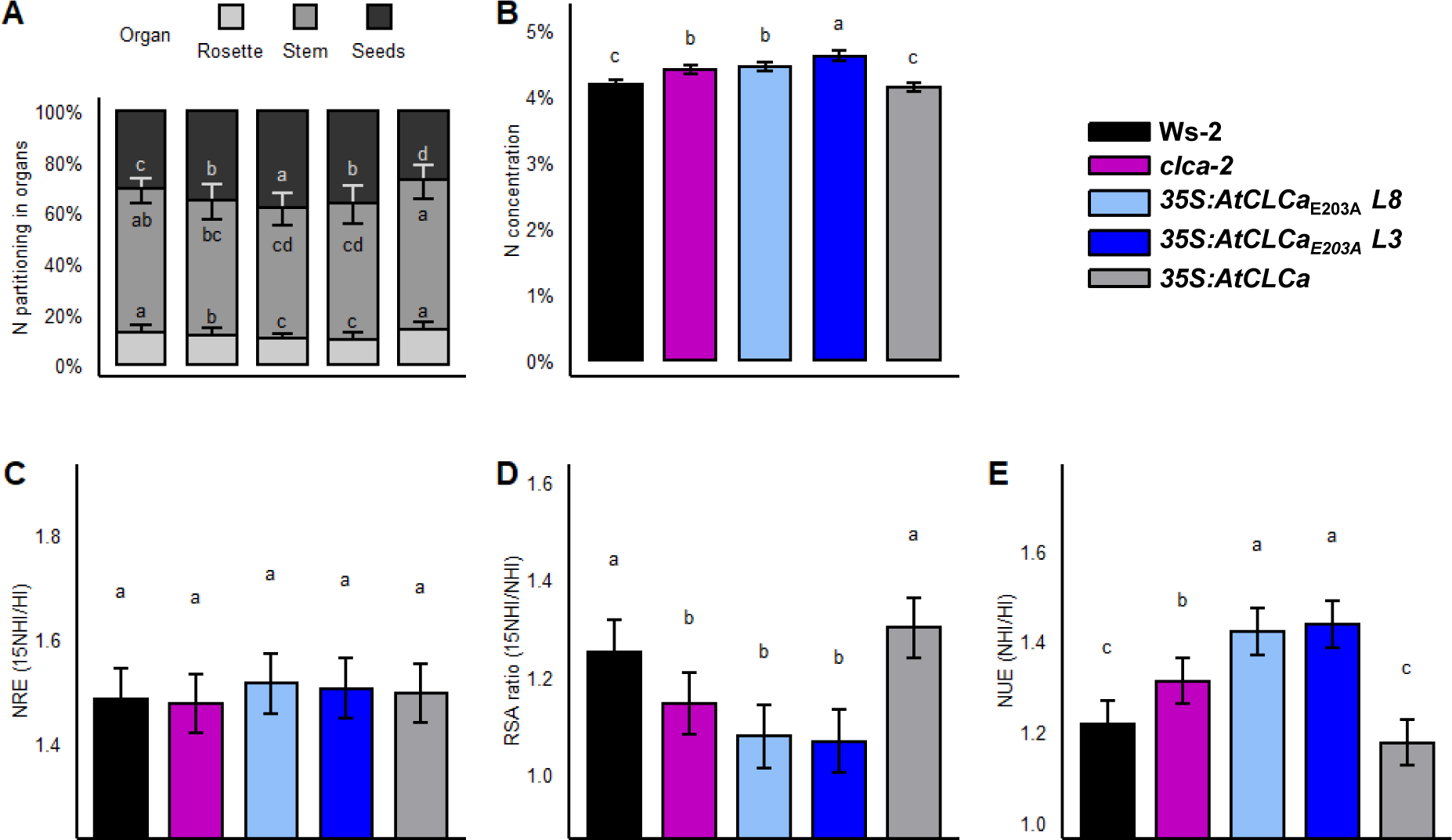
The mutation in the gating glutamate in AtCLCa leads to an increase nitrogen use efficiency at grain-filling stage. Plants of Ws-2, *clca-2*, *clca-2/35S:AtCLCa_E203A_* lines 8 and 3 and *clca-2/35S:AtCLCa* were grown on sand under short days and transferred under long days one week before flowering bud emergence. Plants were harvested at maturity. N partitioning in rosette, stem and seeds **(A)**, seed N concentration **(B)**, N remobilization efficiency **(C)**, the relative specific absorption ratio **(D)** and N used efficiency **(E)**. The results show for each genotype the means ± SD. Statistical analysis was performed using analysis of variance and the means were classified using Tukey HSD test (P<0.05), different letters indicate significant difference.

## DISCUSSION

In eukaryotes, the CLC membrane protein family is formed by both anion/H^+^ exchangers and anion channels. Despite the different transport mode existing in the CLC family the 3D structure of the two kinds of CLCs is surprisingly close (Jentsch and Pusch, 2018). This particular feature has been shown to be mainly related to substitution of the “gating glutamate” in CLC channels. The physiological consequences of the mutations in this residue have been studied in live animals (Novarino et al., 2010; Weinert et al., 2020). So far, the effect of mutating the gating glutamate of CLC exchangers have not been investigated in plants. We used the specific cellular function of AtCLCa in nitrate vacuolar accumulation to investigate the importance of this glutamate residue in plants from the cellular to the whole organism level. We found that the conversion of AtCLCa from an exchanger to a channel, by mutating the “gating glutamate” into an alanine, strongly alters *Arabidopsis* nitrate storage capacity and water homeostasis, leading to reduced plant growth. However, the plants expressing the mutant version of AtCLCa presented a noteworthy increase in N assimilation and NUE compared to wild-type plants.

### The AtCLCa_E203A_ mutation leads to a decrease of nitrate storage in cells

Our results demonstrate that mutating the 203 glutamate of AtCLCa on native vacuolar membrane leads to a partial complementation of the anion current compared to *clca-2* KO mutant (Figure 1). Indeed, AtCLCa_E203A_ is able to transport anions but the currents measured in vacuoles extracted from *clca-2/35S:CLCa_E203A_* plants respond differently to variations of the cytosolic pH compared to plants expressing the wild-type form of AtCLCa. In *clca- 2/35S:CLCa_E203A_*vacuoles, the ionic currents display a reversal potential close to the Nernst potential for NO_3_^-^ confirming that AtCLCa_E203A_ mediates passive ion fluxes independent from the pH gradient existing across the tonoplast. These results confirm the data obtained in a previous work performed in Xenopus oocytes (Bergsdorf et al., 2009). Based on these findings and the seven-fold higher selectivity of AtCLCa for nitrate than for chloride (De Angeli et al., 2006; Wege et al., 2010), we expected that the *clca-2/35S:CLCa_E203A_* lines would be less efficient in accumulating NO_3_^-^ in the vacuole. A passive ion transport system in the tonoplast would drive a vacuolar nitrate accumulation 10-15 times lower than an exchanger (Cookson et al., 2005; De Angeli et al., 2006). In line with this prediction, the kinetic measurements performed *in planta* to analyze nitrate storage and remobilization show a strong decrease of nitrate storage rates in the shoot of *clca-2/35S:AtCLCa_E203A_* compared to Ws-2 and *clca-2* (Figures 5A and 6). Interestingly, in these experiments, we observed an inverse correlation between the accumulation of chloride and nitrate: the plants likely accumulate chloride to compensate the amount of negative charges inside the cells and maintain the electrochemical potential gradients through the membranes when nitrate is scarce (Supplemental Figure S5). These results confirm that AtCLCa_E203A_ mediates passive nitrate fluxes across vacuolar membranes whereas the exchanger activity is not required for efficient accumulation of chloride.

### The coupling of proton and anion transport by AtCLCa is crucial for water content and to ensure the function of nitrate as an osmoticum

Stomata guard cells are widely used as model to study the molecular mechanisms involved in the adjustment of cell turgor. Both *clca-2* and *clca-2/35S:AtCLCa_E203A_* are impaired in stomata aperture in response to light due to an under-accumulation of anions in the vacuoles (Figure 3). This result confirms our previous conclusions on the importance of AtCLCa in the control of the osmotic pressure (Wege et al., 2014). Additionally, comparing wild-type, *clca-2* and *clca-2/35S:AtCLCa_E203A_*plants, we observed here a close correlation between shoot water and nitrate contents (Figures 4A and 5A) which are both reduced in *clca* mutants indicating the importance of AtCLCa in nitrate and water homeostasis. Furthermore, the similar levels of potassium and other main inorganic anions, chloride, phosphate and sulfate, between the genotypes (Figures 5B and 5C) confirm the function of nitrate as an osmoticum (McIntyre, 1997). Our results corroborate previous published work (Cardenas-Navarro et al., 1999). More recently, it was shown, using different mutants of nitrate transporters in Arabidopsis, that the nitrate content in shoots is correlated to the water transport capacity of the roots (Li et al., 2016). This could also be the case in our different genotypes, especially since *AtCLCa* is expressed in mesophyll and guard cells (Geelen et al., 2000; Wege et al., 2014).

In parallel, the characterization of *clca* mutants reveals that AtCLCa and its glutamate 203 are essential to sustain plant fresh weight (Figure 2). Several hypotheses can be proposed to explain this result. First, an altered response of the different transgenic plants to ABA, the hormone involved in water homeostasis, may inhibit the growth. We showed that stomata of *clca-2* and *clca-2/35S:AtCLCa_E203A_* plants respond very weakly to ABA (Figure 3B) which indicates that the sensitivity to ABA is modified in these plants. However, this hypothesis by itself cannot explain the difference observed in cell size and water content between *clca-*2 and *clca-2/35S:AtCLCa_E203A_*lines as the defects in stomata movements are similar in these genotypes. Second, the plants may be disturbed in cytosolic chloride homeostasis, as AtCLCa is able to transport chloride albeit with low affinity compared to nitrate (De Angeli et al., 2006; Wege et al., 2010). The kinetic experiments showed that chloride content does not decrease as much in the *clca-2/35S:AtCLCa_E203A_* lines as in the other genotypes when nitrate is added to the growth medium (Supplemental Figure S5B). Nevertheless, the measured concentrations are below the toxic values and could only account partially for the observed growth phenotype (Jossier et al., 2010). Third, the decrease in stomata aperture may lead to a reduction of gas exchange with the atmosphere and consequently to an inhibition of the photosynthesis rate (Figure 3). In mammals, it has been suggested that the CLC exchangers work in tandem with the V-ATPase to maintain intra-compartment pH maintenance (Satoh et al., 2017). A disruption of pH homeostasis in the different *clca-2* lines could also explain the difference in plant fresh weight (Krebs et al., 2010; Demes et al., 2020). Finally, the simplest hypothesis may be that the decrease of vacuolar NO_3_ ^-^ storage in these genotypes underlies a reduction of cell water potential (Figure 4). It was shown that, during root growth, the vacuolar osmotic potential is important for turgor pressure and to drive cell elongation (Dünser et al., 2019; Kaiser and Scheuring, 2020). It seems plausible that a similar mechanism operates in mesophyll cells: the under-accumulation of nitrate could lead to lower cell expansion which would in turn limit plant growth. Altogether, these results demonstrate the crucial role of a nitrate/proton exchanger on the vacuolar membrane to maintain water homeostasis and cell expansion in Arabidopsis.

### The NO_3_^-^/H^+^ exchanger activity of AtCLCa is essential for the regulation of nitrate assimilation and consequently for NUE

It has been shown that most of the nitrate in leaves is stored in vacuoles of mesophyll cells (Martinoia et al., 1981; Miller and Smith, 2008). The decrease of nitrate in rosette leaves observed in *clca-2/ 35S:AtCLCa_E203A_* plants is most probably due to a reduction of vacuolar nitrate storage. Then this defect in *AtCLCa_E203A_* over-expressing lines probably perturbs nitrate cytosolic homeostasis and, consequently results in the increase of NR activity, which would drive to higher synthesis amounts of amino acids and proteins than in Ws-2 and *clca-2* (Figure 7). These results also confirm that the increase of intracellular nitrate concentration enhances the activity of NR (Aslam et al., 1987). However, the absence of AtCLCa in KO mutant does not lead to increase of amino acids and protein contents as in AtCLCa_E203A_ over- expressing lines, whereas cytosolic nitrate homeostasis is also perturbed in *clca-2* (Monachello et al., 2009). This difference might be explained by the fact that AtCLCa_E103A_ is expressed under the 35S promoter and is most likely active in cells of these transgenic lines in which it is not normally expressed in wild-type plants.

Among the amino acids over-accumulated in the *clca-2/35S:AtCLCa_E203A_*transgenic lines (Figure 7B), asparagine and glutamine are amongst the ones preferentially transported through the plant (Havé et al., 2017). Serine and glycine, also accumulated in the over-expressors, may reflect an increase of photorespiration, the pathway that supplies reductants such as NADPH necessary for nitrate assimilation in C3 plants (Migge et al., 2000; Oliveira et al., 2002; Bloom, 2015). Analyses of the activity of this metabolic pathway will have to be performed to confirm this hypothesis. In parallel, *clca-2/ 35S:AtCLCa_E203A_*lines accumulate four time more malate in than wild-type (Figure 5D). This increase is not trivial, as malate is a trivalent anion that could compensate nitrate negative charge depletion charges in the vacuolar lumen when nitrate concentration is decreased. However, malate is also a substrate of the photorespiratory pathway producing reductants. Further, the NR activity requires two protons to reduce one molecule of nitrate into nitrite (Feng et al., 2020). Malate synthesis could then be stimulated to compensate the alkalization due to the increase of nitrate assimilation (Eisenhut et al., 2019; Bloom, 2015; Feng et al., 2020).

At the reproductive stage, *clca-2* and, to a greater extent, *clca-2/35S:AtCLCa_E203A_* lines showed a different allocation of nitrogen in the plant organs, resulting in a higher NUE (Figure 8). These results reflect an enhanced ability of seeds to store N independently of the plant to produce seeds and indicate variation in N fluxes during the reproductive phase. Expression of *AtCLCa* was reported in roots, rosette leaves and siliques, but not in seeds (David et al., 2014; Geelen et al., 2000). The large increase in seed N allocation in *clca-2/ 35S:AtCLCa_E203A_* lines compared Ws-2 and *clca-2* probably results from an indirect effect of AtCLCa defect in the rosette and stem compartments (stem inflorescences + silique envelopes). Surprisingly, the change in NUE was found not to be due to an increase in N remobilization under this growth condition as NRE values were similar between the various lines. In contrast, the RSA ratio was significantly reduced in the *clca-2* background lines indicating a higher N uptake after flowering. Our previous study showed that root nitrate influx is reduced in *clca* KO mutants but measurements were performed at vegetative stage (Monachello et al., 2009). However, more recent work suggested that vacuolar nitrate transporters like CLCs drive an increase in root N uptake and/or a higher root/shoot translocation at later stages (He et al., 2017; Li et al., 2020) and may explain the low values of RSA ratio obtained in *clca-2* lines in our study. The difference of NUE observed between *clca-2* and *clca-2/35S:AtCLCa_E203A_* lines could be due to a lower nitrate storage capacity in both the rosette and reproductive organs in *AtCLCa_E203A_* over-expressors. This result illustrates the importance of nitrogen storage in leaves, stem inflorescence, and siliques for nitrate uptake by roots and nitrogen allocation in plant organs during seed filling. Nitrate uptake relies on NO_3_^-^/H^+^ symporters. The changes in cytosolic nitrate concentration, cytosolic pH and nitrate assimilation linked to the absence of AtCLCa or the presence of the uncoupled form of AtCLCa could then modify the activities of these plasma membrane transporters (Feng et al., 2020; Filleur and Daniel-Vedele, 1999). Altogether, this finding highlights the importance of the proton antiport activity of AtCLCa for regulating nitrate assimilation and consequently NUE.

## Conclusion

Based on peptide sequence analysis of the closest homologs of AtCLCa from algae, lycophytes, bryophytes and spermaphytes, each species has conserved at least one CLC with the gating glutamate residue (Supplemental Figure S8). This conservation suggested a very strong importance of this residue in the green lineage. Our study on the physiological consequences of the mutation in the gating glutamate of AtCLCa provide insights on the selective pressure underlying the conservation of an exchanger rather than a channel for this protein. Although this mutation leads to a higher plant nutritional value and better NUE, it also induces a decrease of plant growth due to water homeostasis disruption expected to severely impair plant fitness. The conservation of the exchange mechanism of AtCLCa is then likely to be correlated to the maintenance of water homeostasis irrespective of the external nitrogen fluctuations. A previous study showed that a decrease of the nitrate vacuolar sequestration in roots induces a higher translocation to the shoot, a higher assimilation and biomass (Han et al., 2016). Therefore, we could wonder if a root-specific expression of *AtCLCa_E203A_* would provide plants with high protein level and NUE, but without the associated growth disruption probably due to the expression of *AtCLCa* in mesophyll and guard cells. This may give new clues to generate plants with higher NUE without disturbing water homeostasis.

## MATERIALS AND METHODS

### Accession number

Sequence data from this article can be found in the GenBank/EMBL data libraries under the accession number At5g40890.

### Plant material

Experiments were performed on *Arabidopsis thaliana* (accession Wassilewskija [Ws-2]) wild-type plants and T-DNA insertion mutant *clca-2* (De Angeli et al., 2006). The *clca- 2/35S:AtCLCa* complemented line was produced in a previous work (Wege et al., 2010). AtCLCa_E203A_ point mutation was introduced in *AtCLCa* cDNA using the QuikChange II XL Site-Directed Mutagenesis Kit (Stratagene) into the Gateway vector pH2GW7.0 (Karimi et al., 2002) under the control of the 35S promoter or into the Gateway pMDC43 vector (Curtis and Grossniklaus, 2003) allowing the fusion of GFP at the N-terminal part of *AtCLCa*. For *clca-2/pAtCLCa:AtCLCa* and *clca-2/pAtCLCa:AtCLCa_E203A_* lines generation, a 1.9-kb fragment of *AtCLCa* promoter was produced by PCR amplification on genomic DNA ([Ws-2] accession) using purified primer pair: 5’- nnnnncccgggggttttgccactcatacttt-3’ (Forward) and 5’-nnnnnactagttgggtggatgggtaccatat-3’ (Reverse). The PCR fragment was cloned into pH2GW7.0 between *SmaI* and *SpeI* restriction sites upstream *AtCLCa* or *AtCLCa_E203A_* cDNA sequences. Those constructs were used to transform T-DNA knockout plant for *AtCLCa* (*clca-2*) by floral-dipping (Clough and Bent, 1998). The seeds were selected on hygromycine B (20 µg.mL^-1^) and two T3 homozygous lines were chosen.

### Plant growth conditions

All experiments were performed on plants grown under short days conditions (8h light, 16h dark) at 22°C, 60% relative humidity, 75 µE light intensity. Water and potassium content experiments were performed in plants grown for five to seven weeks in Jiffy® peat pellets. For fresh weight and anion, amino acids and proteins content determinations, plants were grown hydroponically for four to five weeks. Seeds were sterilized and sown on seed-holders (Araponics, Liège, Belgium) filled with half-strength MS medium containing 0.60% phytoagar. The boxes were filled with MilliQ water, put at 4°C for 4 days for seeds stratification, and then transferred in the culture room. Once roots have emerged in water solution, the medium is replaced by a modified Hoagland nutrient solution (1.5 mM Ca(NO_3_)_2_, 1.25 mM KNO_3_, 0.75 mM MgSO_4_, 0.28 mM KH_2_PO_4_, micronutrients [KCl 50 µM, H_3_BO_3_ 25 µM, ZnSO_4_ 1 µM, CuSO_4_ 0.5 µM, Na_2_MoO_4_ 0.1 µM, MnSO_4_ 5 µM], chelated iron Fe-HBED 20 µM and MES 2 mM pH 5.7 with KOH). Nutrient solutions were replaced twice a week. For nitrate starvation experiments, five weeks plants roots grown in hydroponic system were rinsed twice in the nitrate starvation medium (Ca(NO_3_)_2_ replaced by CaSO_4_ 1.5 mM and KNO_3_ by KCl 1.25 mM) and plants were put on the starvation medium for 120h. For nitrate storage experiment, 4 weeks plants were nitrate starved for 10 days, then nitrate starvation medium was replaced by complete Hoagland nutrient solution for 120 h. For all of these experiments, six plants per genotype for each time point were harvested, rosettes and roots separately. Those experiments were performed three times.

For measurement of NUE at reproductive stage, seeds were stratified in tubes containing water in a cold room for 2 days at 4°C in the dark. After stratification, seeds were directly sown on sand and watered with a 10-mM nitrate solution. Plants were grown in a climatic chamber, in short days (8/16 h day/night photoperiod) during 7 weeks then transferred in long days (12/12 h day/night photoperiod) until final harvest. Composition of nutritive solution is described in Chardon et al. (2010). Six to nine plants per genotype were harvested. The experiment was performed twice.

### 15N labelling and determination of N partitioning, N remobilization, relative specific absorption and NUE

On week before the transfer, when the plant were in exponential vegetative growth, plants were watered with a 10 mM ^15^NO_3_^-^ 10% enrichment solution. To analyze unlabeled samples, a few ^15^NO_3_^-^-free plants were harvested in order to determine the ^15^N natural abundance. After two days, sand was rinsed twice in osmotic water baths. At the end of their cycle, when all seeds were mature and the rosette dry, plants were harvested. Samples were separated as (i) rosette (rosette leaves), (ii) stem (inflorescence stem + cauline leaves + empty dry siliques), and (iii) seeds (total seeds). The dry weight of rosette, stem and seeds was determined. Subsamples of 1000–2000 μg were carefully weighed in tin capsules to determine the total N percentage (N% as mg (100 mg DW)^−1^) and the ^15^N abundance using a FLASH 2000 Organic Elemental Analyzer (Thermo Fisher Scientific) coupled to a Delta V Advantage isotope ratio mass spectrometer (Thermo Fisher Scientific). The ^15^N abundance in each sample was measured as atom percent and defined as A%=100×(^15^N)/(^15^N+ ^14^N). In unlabeled plant controls, A% control was 0.3660. The ^15^N enrichment (E%) of the plant material was then calculated as (A%sample–A%control). The absolute quantity of N and ^15^N contained in the sample was calculated as QtyN=DW×N% and Qty^15^N= DW×E%×N%, respectively. Different parameters used to evaluate HI, NUE, N remobilization, and its components were defined as follows:

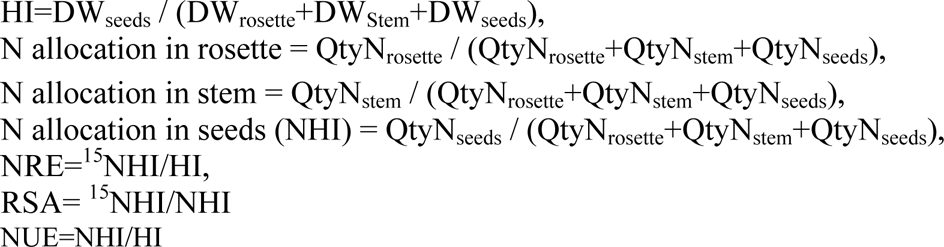

### RT-qPCR analysis

Total plants RNA were extracted from 4 weeks plants using the RNeasy kit (Qiagen, Germany) and two micrograms of RNA was reverse-transcribed using SuperScript IV^TM^ Reverse Transcriptase according to the manufacturer’s instructions (Thermo Fisher Scientific). Real-time PCR was performed on cDNAs in a final volume of 10 µL using SYBR Green I master mix (Roche Life Science) and primers for *AtCLCa* gene, 5’- atcaaatggagatggcttcg-3’ (Forward) and 5’-cctcaagagcgaaaagtactc-3’ (Reverse), and *Actin2* reference gene, 5’-ggtaacattgtgctcagtggtgg-3’ (Forward) and 5’-aacgaccttaatcttcatgct-3’ (Reverse). The reactions were performed in a LightCycler^®^ 96 Real-Time PCR system (Roche Life Science). Samples were subjected to ten minutes of pre-incubation at 95°C, then 45 amplification cycles with 15 seconds at 95°C, 15 seconds at 60°C and 15 seconds at 72°C. A high resolution melting was performed to assess amplification specificity and several cDNA dilutions were tested to perform primers efficiency calculation. The results were analyzed using the LightCycler^®^ Software (Roche Life Science) and normalized with the *Actin 2* gene expression.

### Confocal microscopy

AtCLCa_E203A_ localization was checked with a confocal microscope Leica TCS SP8 using the fusion of GFP at the N-terminal part of AtCLCa. The GFP was excited at 488 nm and its fluorescence emission signal was analyzed between 500 and 525 nm.

### Electrophysiological experiments

Vacuoles for electrophysiological experiments were extracted from *A. thaliana* mesophyll protoplasts as described before (Song et al., 2003). Patch clamp recordings were performed in the whole-vacuole confirmation and recorded with a HEKA amplifier EPC-10 USB (HEKA, Lambrecht-Pfalz, Germany). The recordings were acquired and controlled with the software PatchMaster (HEKA, Lambrecht-Pfalz, Germany). Currents were induced by five-second pulses from -97 to +63 mV in 20 mV increments and the potentials were corrected by the liquid junction potential (Neher, 1992). Standard solutions contained: (vacuolar) 200 mM BTP NO ^-^, 2 mM MgCl_2_, 0.1 mM CaCl_2_, 10mM MES pH 5.5; (cytosolic) 20 mM Bis-Tris-Propane (BTP) Cl^-^, 10 mM MES pH 7, 0.1 mM CaCl_2_, 2 mM MgCl_2_; pH 7. Cytosolic NO_3_^-^- solutions containing 0.1 mM Ca(NO_3_)_2_, 2 mM Mg(NO_3_)_2,_ 10 mM BTP were adjusted to pH 7 or pH 9 with MES. The osmolarity of the solutions was adjusted to π= 600 mOsm with sorbitol. Only measurements of stable vacuoles that returned to initial reversal potentials in the starting conditions were considered.

### Plants compounds content and nitrate reductase activity measurements

For nitrate and chloride content measurements, shoot and root of five weeks old plants were harvested separately, weighted and fast frozen in liquid nitrogen. Plants material was grounded, homogenized in 1 mL (shoot) or 500 µL (root) of MilliQ water and exposed to three successive freeze-thaw cycles. After the last thawing, plants material was centrifuged 10 minutes at full speed to pellet cell debris and recover the supernatant that will be used for nitrate and chloride colorimetric assays (Miranda et al., 2001). For anion content determination, the shoot samples were used for HPLC analysis (ICS5000, ThermoFisher) or for malate determination using the Biovision^TM^ kit. For potassium content analysis, shoot were harvested and dried at 60°C 3 days. The dried samples were digested into 2 mL of 70 % nitric acid in a DigiBlock ED36 (LabTech) at 80°C for 1 h, 100°C for 1 h and then 120°C for 2h. After dilution in ultra-pure water, the potassium content was determined by atomic absorption spectrometry using an AA240FS flame spectrometer (Agilent Technologies).

The nitrate reductase activity was determined as described in Kim and Seo (2018). For amino-acids quantification, plants shoots were fast frozen in liquid nitrogen and lyophilized overnight. Dried material was weighted to equalize the amount of samples and finely grounded in liquid nitrogen with a pestle and mortar. Polar metabolites were extracted into 80% methanol, 20% water containing 0.2 mM α -Amino-n-Butyric Acid (α-ABA) as an internal standard. The samples were centrifuged and several aliquots were dried overnight under vacuum. Prior to HPLC analysis (Waters Alliance instrument with a Waters 2475 multi-wavelength fluorescence detector), aliquots were resuspended in milliQ water and filtered into autosampler vials before precolumn derivatization. Standard amino-acids solutions were used for calibration curve generation; a correction was performed using internal standard variation and normalized with dry weight. For soluble protein content determination, 100 mg of each plant was harvested and grounded in liquid nitrogen. Then, 350 µl of extraction buffer (50 mM Hepes/NaOH pH 7.2, 1.5 mM MgCl_2_, 1 mM EGTA 0.2M, 10% glycerol, 1% Triton, 2 mM PMSF, 150 mM NaCl, antiprotease-EDTA) were added before vortex homogenization. Samples were incubated under agitation for 30 minutes at 4°C. After centrifugation, supernatant were recovered and used for total soluble protein content determination using a standard Bradford assay (Bradford, 1976).

### Stomata dynamics measurements

Stomata bioassay were performed on five weeks-old plants as described in Jossier et al. (2010). Two hours before the beginning of the light period, plants were collected and two leaves per genotype were glued on cover slides with surgical glue (Hollister Medical Adhesive, Adapt^TM^ 7730) to peel the epidermis. Those cover slides were immediately immerged in MES/KCl buffer (50 mM, pH 6.15 with KOH) and kept in the dark for 1 hour. After this dark period, images were acquired for initial stomata aperture measurements and then the aperture was monitored after opening induction by 75 µE of light for 4h30 at 22°C. For stomata closing experiment, additionally, the epidermis were incubated for 3 hours with 50 µM of Abcissic acid and the aperture area was determined. Images acquisition was performed with a 40X objective on a wide field inverted microscope (DMI600B, Leica, Imagerie Gif, Gif-Sur-Yvette) coupled with a Hamamatsu camera. To capture stomata images, Z-stacks were acquired to obtain a clear image of all cells. Ostiole area determination was performed automatically on different types of z-projections, in a procedure we developed with ImageJ software (Schneider et al., 2012); Supplemental methods). Measurements were performed on 86-150 stomata per genotype per treatment (two leaves) and repeated three times.

### Plants water content measurements

For water content analysis, plants rosettes were harvested and weighted (n=10 for each genotype, N=3 biological replicates), rosettes were dried for three days at 65°C and relative water content was calculated as : (FreshWeight-DryWeight)/FreshWeight. Relative water contents were determined similarly during the dehydration tests performed under a laminar flow hood on seven weeks plants as described by Wege et al. (2014).

### Cell size determination

Flow cytometry was used to determine relative cell size of plants leaves. On five weeks old plants, seven leaves of three plants per genotype were harvested and digested with an enzymatic mix (1 % cellulase R-10, 0.2 % Macerozyme R-10, 0.4 M Mannitol, 20 mM KCl, 20 mM MES/KOH pH 5.7, 10 mM CaCl_2_, 0.1% w/v BSA) for three hours. Protoplasts were retrieved by centrifugation at low speed (100 g) for two minutes. Protoplasts were re- suspended in an appropriate solution (1 mM CaCl_2_, 10 mM MES pH 5.3 (KOH), 594 mOsm with sorbitol). Then, they were filtered through a 50 µm nylon filter, and analysed on a MoFlo Astrios cytometer, driven by Summit 6.3 (Beckman-Coulter). Chlorophyll was excited by a 488 nm solid-state laser (150 mW), taking emission at 664/22 nm. Forward Scatter (FSC, size) and Side Scatter (SSC, granularity) were taken on the 488 nm laser. The first region of interest (gate) was focused on events with high homogenous fluorescence in chlorophyll. Then, mean values of FSC-Area and SSC-Area parameters were taken with the same gating strategy for each sample. Each histogram comprised more than 30,000 protoplasts.

## AKNOWLEDGMENTS

We thank Romain Le Bars (Imagerie-Gif Platform, I2BC, Gif-sur-Yvette, France) for his technical support with confocal microscopy, Caroline Mauve et Françoise Gilard (Metabolomic platform, IPS2, Gif-sur-Yvette, France) for their help analyzing amino acids contents. Thank to Eugene Diatloff for its critical reading of the manuscript. This work has benefited from Imagerie-Gif core facility supported by the ‘Agence Nationale de la Recherche’ (ANR-11-EQPX-0029/Morphoscope, ANR-10-INBS-04/FranceBioImaging; ANR-11-IDEX-0003-02/ Saclay Plant Sciences), is supported by an ANR grant (ANR- VACTION: ANR-16-CE92-0004-21), the LabEx Saclay Plant Sciences-SPS (ANR-10- LABX-0040-SPS), the ‘Centre National de la Recherche Scientifique’ (CNRS), the University Paris Cité and the University Paris-Saclay.

## AUTHOR CONTRIBUTIONS

A.D.A., S.T. and S.F. designed the project. J.H., C.L., C.E. and F.A.C. performed the experiments, M.B. conducted the cytometry measurements. M.W.B. developed the macro running under Image J to analyze the ostiole area. J.H., C.L., C.E. and A.D.A. analyzed the data. A.M. and F.C. performed and analyzed the NUE experiment. J.H., C.L. and S.F. wrote the manuscript. All authors revised the article.

